# Predicting Collaborative Learning Quality through Physiological Synchrony Recorded by Wearable Biosensors

**DOI:** 10.1101/2020.06.01.127449

**Authors:** Yang Liu, Tingting Wang, Kun Wang, Yu Zhang

## Abstract

Interpersonal physiological synchrony has been consistently found during collaborative tasks. However, few studies have applied synchrony to predict collaborative learning quality in real classroom. This study collected electrodermal activity (EDA) and heart rate (HR) in naturalistic class sessions, and compared the physiological synchrony between independent task and group discussion task. Since each student learn differently and not everyone prefers collaborative learning, participants were sorted into collaboration and independent dyads based on collaborative behaviors before data analysis. The result showed that during groups discussions, high collaboration pairs produced significantly higher synchrony than low collaboration dyads (*p* = 0.010). Given the equivalent engagement level during independent and collaborative tasks, the difference of physiological synchrony between high and low collaboration dyads was triggered by collaboration quality. Building upon this result, the classification analysis was conducted, indicating that EDA synchrony can predict collaboration quality (AUC = 0.767, *p* = 0.015).

## Introduction and Related Literatures

In a world that is deeply connected, collaborative learning is believed to be the most important way of learning, shared knowledge construction, decision making, critical thinking, and problem solving (Bruffee, 1999; Dillenbourg, 1999; Gokhale, 2012; Van Kleef, De Dreu, & Manstead, 2010). Scholars and practitioners advocating collaborative learning believe that learning is inherently active, constructive, and social. Successful collaboration benefits the whole group by immersing the students in an active learning condition to increase engagement and joint attention, relearn through retrieval, negotiate multiple perspectives, increase working memory resources, to name a few (Barron, 2003; Johnson & Johnson, 1985; Kirschner, Paas & Kirschner, 2009; Kuhn & Crowell, 2011; Roediger & Karpicke, 2006). Broader education goals such as involvement, cooperation and teamwork, and civic responsibility are also believed to be achieved by collaborative learning (Smith & Macgregor, 1992).

Dillenbourg (1999) provided a general definition of collaborative learning as “a situation in which two or more people learn or attempt to learn something together.” In this definition, “learn something” was broadly interpreted as activities including “follow a course”, “study course material”, “perform learning activities such as problem solving”, “learn from lifelong work practice”; and “together” was interpreted as different forms of interaction. In fact, individual interaction is crucial in successful collaborative learning (Hiltz & Turoff, 2002; Kreijns, Kirschner, & Jochems, 2003; Soller, Goodman, Linton, & Gaimari, 1998), and thus serves as a key component of collaboration quality. As quite a few factors may affect the quality of individual interaction, emotion is one of the most significant and emotions moderate human behaviors in observable patterns (Balters & Steinert, 2017). Emotion regulation abilities is highly related to the success of interpersonal interactions, especially when the individuals collaborate with peers or in the workplace (Eligio, Ainsworth, & Crook, 2012; Lopes, Salovey, Côté, Beers, & Petty, 2005).

Given the importance of collaborative learning, the measurement of its quality is, however, very complex and challenging. Existing approaches can be categorized into subjective and objective measurements. Subjective measurement mainly relies on self-report data, including both interview (Salovaara & Järvelä, 2003; Sidani & Reesem, 2018) and scales (Orchard, King, Khalili, & Bezzina, 2012; OECD, 2013); while objective measurement mainly relies on explicit observational data, which mainly captures verbal communication and non-verbal behaviors (Chi et al., 2018; Marty & Carron, 2011; Mehl, 2017; Odom & Ogawa, 1992). The main defect of self-report data is the subject perspective that could be manipulated by the participants. For instance, the social desirability bias is a famous potential threat (Fisher, 1993; Holbrook & Krosnick, 2009).

Analyzing the verbal content of interactions is the most straightforward approach for analyzing the quality of interaction, in both face-to-face and computer supported contexts and has been commonly used in educational and psychological studies (Grau & Whitebread, 2012; Kent, Laslo, & Rafaeli, 2016; Vuopala, Hyvönen, & Järvelä, 2016). The limitations of content analysis are, however, labor intensive and difficult to provide instant feedback to either students or teachers. Non-verbal interactions such as eye contacts, facial expression, and body movement are also possible to be captured and analyzed (Montague, Chen, Xu, Chewning, & Barrett, 2013; Tunçgenç & Cohen, 2018), thanks to the rapid development of computing capability and machine learning algorithm. These methodologies have shown rich effectiveness in the interpretation of human behaviors (Elo & Kyngäs, 2008; Hsieh & Shannon, 2005). However, there are still issues on trustworthiness of both the content analysis and the interpretation of explicit behaviors (Elo, et al., 2014). The implicit factors such as emotional contagion and affect infusion among individuals, on the other hand, may be more crucial to cognition (Okon-Singer, Hendler, Pessoa, & Shackman, 2015), but is far from being fully investigated due to challenges in measurement and data constraint (Fujiki, Brinton, & Clarke, 2002).

Emotion plays an important role in education, especially in collaborative tasks (Järvenoja & Järvelä, 2009; Schutz & Pekrun, 2007). The effect can be either positive or negative (Imai, 2010). Comparing to verbal content, emotional state is directly detectable though quantitative measurements. Building upon the theory on human automatic nerves system (ANS), important components of collaborative learning, such as cognitive load and emotional state, are believed to be monitorable through neurophysiological signals, including EEG, fNIRs, ECG, and EDA. Arousal and valence can be evoked and detected in specific situations (Agrafioti, Hatzinakos, & Anderson, 2010; Boucsein, 2012; Dawson, Schell, & Filion, 2017; Ramirez & Vamvakousis, 2012). The effects of individual interactions on emotion can also be measured through multimodal physiological signals (Heaphy & Dutton, 2008; Mønster, Håkonsson, Eskildsen, & Wallot, 2016).

Neurophysiological signals have been considered as promising measurements of emotional characteristics and can capture students’ learning process that go beyond acquisition of knowledge (Léger, Davis, Cronan, & Perret, 2014; Ochoa & Worsley, 2016). Positive evidences on the correlation between interpersonal neurophysiological synchrony and interaction are consistently reported in recent years. Using various hyperscanning technologies, inter-brain synchrony has been identified during face-to-face communication or interactive decision making (Dikker et al., 2017; Hu et al., 2018; Jiang et al., 2012), suggesting special neural processes recruited by interaction. Interpersonal physiological synchrony has also been consistently found during collaborative tasks and used as indicators of effective collaboration. Higher level of synchrony is associated with better task performance and learning gains in collaborative tasks (Ahonen et al., 2016; Dich, Reilly, & Schneider, 2018; Pijeira-Díaz, Drachsler, Järvelä, & Kirschner, 2016).

However, brain hyperscanning devices are usually not easily to use in real classroom learning, because of people’s concern on health safety issues of brain hyperscanning, and their visual interruption to both students and teachers. Data quality might also be a problem in naturalistic settings (Matusz, Dikker, Huth, & Perrodin, 2019). Physiological signal, on the other hand, can be easily and steadily recorded at distal sites such as fingers and wrists (Boucsein, 2012) and thus easily accepted by parents and students. The majority of existing studies that measure physiological synchrony in collaborative learning are basically lab-based experiments (Dich et al., 2018; Pijeira-Díaz et al., 2016; Dindar, Alikhani, Malmberg, Järvelä, & Seppänen,2019). The tasks include open-ending problem-based learning topics such as designing breakfast for marathoners (Haataja, Malmberg, & Järvelä, 2018), or pair-programming task with restricted solutions (Xie, Reilly, Dich, & Schneider, 2018). Although a few of them collected data in real classroom (Ahonen, Cowley, Hellas, & Puolamäki, 2018), they only reported correlation between synchrony and collaboration, but did not further explore the practical potential of predicting collaboration through synchrony in naturalistic scenarios.

Naturalistic scenario, instead of laboratory setting, is crucial in the research of collaborative learning. First, it is social in nature and cannot be simulated in fully controlled, isolated environment. Second, it is constructive and targets at high-level cognitive skills such as problem-solving, knowledge construction, and collaboration, and cannot be substituted by simple cognitive tasks, which are frequently used in laboratory investigations. Third, even applying ethologically relevant stimuli in a laboratory context, participants may react differently in both behaviors and neurocognitive signals (de Heer, Huth, Griffiths, Gallant, & Theunissen, 2017; Qu et al., 2020). In the real classroom context, the interaction behavior varies across students, which is very different from laboratory settings, where participants will try their best to comply to the research design. In fact, naturalistic real-world research is believed to be necessary to understanding human behaviors in neuroscience (Matusz et al., 2019). Classroom learning can serve as an ideal semi-structured scenario to bridge the laboratory based research and the real world.

The rapid development of wearable biosensing technologies makes it possible to record neurophysiological signals during naturalistic classroom learning. Recently, researchers used portable EDA and EEG sensors in classroom to record the neurophysiological signals of teachers and students in variance of learning conditions, including lectures, discussion, movie viewing and real exam (Dikker et al., 2017; Poulsen, Kamronn, Dmochowski, Parra, & Hansen, 2017; Zhang et al., 2018; Qu et al., 2020). By recording physiological signal in fully real-world collaborative learning, the present study attempts to apply physiological synchrony to predict interaction quality in real collaborative learning. Although collaborative activities can range from classroom discussions to team research that covers a whole semester or year, this study focuses on classroom discussion as it is the simplest and most general scenario among diverse collaborative learning approaches.

In the current study, the researchers collected physiological and behavioral data from naturalistic class sessions and analyzed interpersonal synchrony during individual and collaborative tasks. Students were naturally divided into two kinds of collaborative dyads according to their different learning styles as captured by behaviors, i.e., collaborative and independent dyads. The result showed a significant difference in interpersonal physiological synchrony between collaborative dyads and independent dyads, i.e., high and low interactive levels. Following classification analysis confirmed the potential of applying physiological synchrony as an indicator of collaboration quality. This finding is promising in future applications of evaluation in student learning style.

## Method

### Participants

Participants were recruited from an undergraduate level elective course that requires no prerequisite domain knowledge. The sixteen students who registered in this course were from twelve different departments and programs across natural sciences, pharmaceutical science, engineering, social sciences, and humanities. Data collection last for two class sessions in two consecutive weeks. Fifteen out of sixteen students (M = 21.61, SD = 2.43, 8 females) signed the informed consent form at the beginning of the first data collection session. One student quit at the second session due to health conditions.

Sixteen students formed four 3-people and one 4-people discussion groups at the beginning of the semester, resulted in 16 dyads in the first data collection session. In the second session, there were 14 dyads since one person quit from a 3-people group, making a total of 30 dyads. Dyad No. 15 was eliminated from all analyses and No. 13 and No. 15 were eliminated from analyses on collaborative learning because one student in this group shared laptop screen with their teammate and discussed (pair No. 15) during the independent task, violating the experiment requirement (Figure 1).

**Figure 1.**
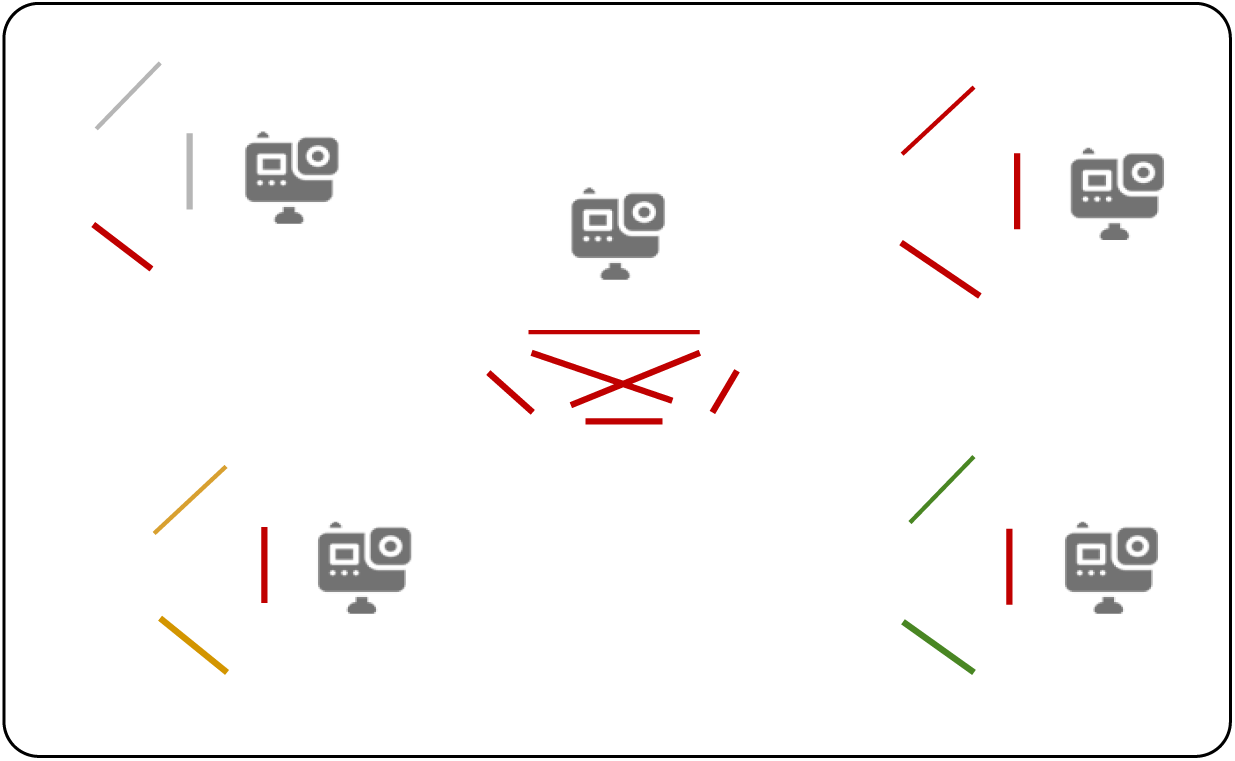
Classroom setting and intra-group dyads. The grey figure did not participate in the study at all. The yellow figure and nodes were excluded from the first data collection session because of violation of experiment requirement. The green figure quit the study in the second-round data collection session because of health conditions. Thus, a total of 28 dyads were used in the analyses.

### Experimental Tasks and Materials

In the second half of each class session, after the lecture, the instructor assigned an open-ended problem to the class. Students were required to solve the problem independently first (independent task, IT), followed immediately by a group collaborative discussion (interaction analyzed in pairs, PT) on the same problem. The problems in both steps were the same, except that participants were asked to solve the problem alone or with group members. In the first session, the students were asked to review course materials and sort out a list of key knowledge by its significance, then discuss with their group members to forge a comprehensive agreement on the list. In the second class session, the students were asked to explore new approaches for engagement measurements alone and then discuss with their group members to form a comprehensive approach.

A short survey was used to evaluate participants’ engagement level and emotional state during IT and PT respectively. Engagement was measured with a five-point Likert scale. The emotional state was measured with a five-scale Self-Assessment Manikin (SAM) to rate the affective dimensions of valence, arousal, and dominance (Bradley & Lang, 1994).

### Settings and Apparatus

The settings followed the naturalistic class settings of this course. Each student had their own chair desk with rolling wheels. Skin conductance (GSR) and heart rate (HR) was collected from each participant using the unobtrusive Huixin Psychorus wristband, capturing data at a sampling rate of 40Hz for GSR and 1Hz for HR. Each group was videotaped during both independent task and collaborative discussion.

### Procedures

Before the beginning of the first data collection session, researchers collected the signed informed consent and helped the students wear the wristbands properly to ensure good data quality.

There was a 3-min close-eye baseline session and a 2-min open-eye baseline session before the independent task. After the baseline sessions, all instructions were given by the instructor. The independent step last for 7-10 minutes and the group collaborative learning task last for 12-17 minutes. The short survey was collected immediately after IT and PT (Figure 2). Two to five minutes were cut from the beginning of the PT sessions to eliminate any continued effect from the IT sessions in data analysis. Students’ own physiological data reports were provided to them after the data collection to appreciate their participation.

**Figure 2.**
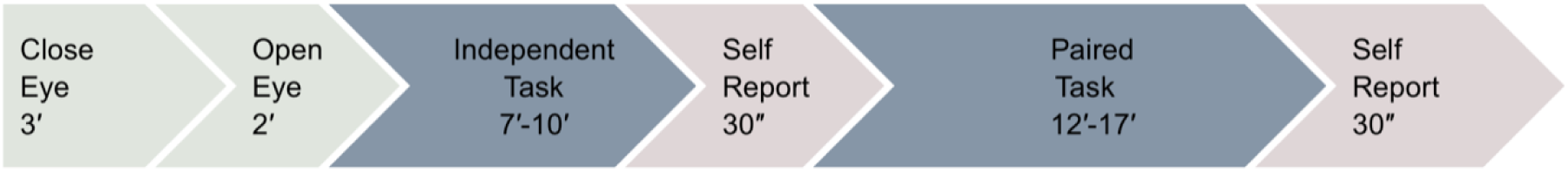
Procedure of the class data collection.

### Physiological Data (GSR) Preprocessing

The visual inspection was performed to control the quality of the raw skin conductance (GSR) signals. Since the number and magnitude of the artifacts were not significant, manual artifacts removal was skipped to minimize the influence on the raw data. The signals were then smoothed using the Gaussian smoothing algorithm and was down sampled from 40Hz to 10Hz in a MATLAB-based skin conductance analysis software (Ledalab 3.4.9; www.ledalab.de; Benedek & Kaernbach, 2010).

## Analysis and Results

### Ground truth and procedure of analysis

In naturalistic setting without artificial design, ground truth should first be defined before physiological data analysis. The ground truth is that students learn differently and not everyone prefers/suites collaborative learning. In the real class sessions, students acted in their own learning style, which refers to the stable trait which decides how learners perceive and respond to learning environments (Keefe, 1979). The Felder and Silverman mode categorizes students’ learning style into five dimensions: active/reflective; sensing/intuitive; visual/verbal; sequential/global; and inductive/deductive (Alfonseca et al., 2006; Felder & Silverman, 1988). Active learners prefer to internalize information from external environments, and they are more likely to share opinions with peers frequently during collaboration; however, reflective learners tend to examine and process information by themselves (Felder & Silverman, 1988).That is, even though the students were equally engaged in the tasks, they may generate different perspectives and preferences for collaborative learning mode, and use different cognitive strategies during interaction (Cabrera et al., 1998; Kayes, 2005). The homogeneity of learning styles within a group will also affect the interactive effect, and the higher homogeneity group members may have better collaboration quality (Alfonseca et al., 2006).

Thus, participants were sorted into collaborative and independent dyads based on their collaborative behaviors. Collaborative dyads are expected to have higher quality interaction during collaborative task and generate higher physiological synchrony.

A series of hypothesis tests were conducted using mean comparisons along different dimensions. Classification analysis was then applied to see if physiological synchrony can predict collaborative learning quality. Figure 3 indicates the logic of the whole analysis.

**Figure 3.**
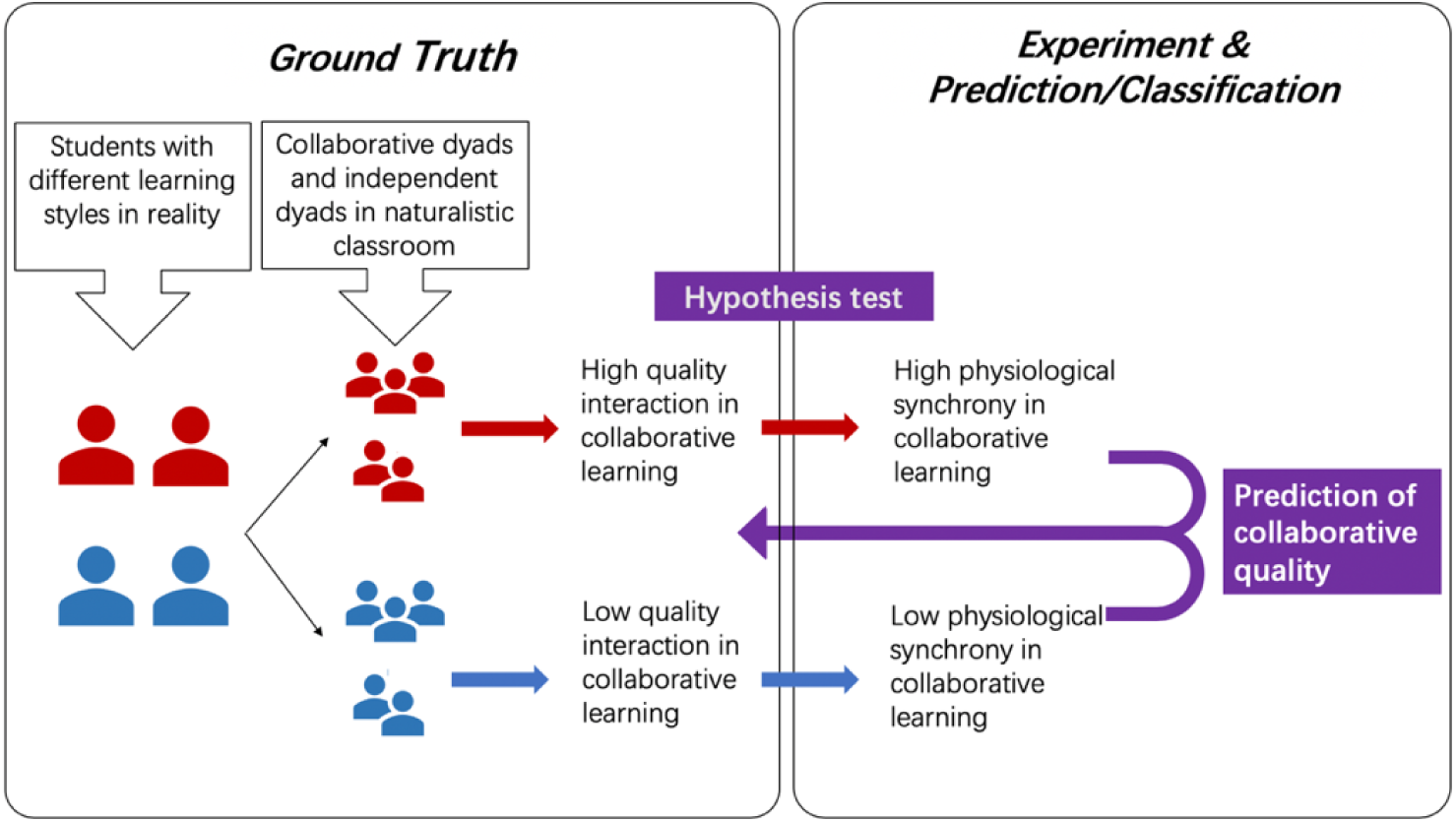
Participants were categorized into collaborative dyads (in red) and independent dyads (in blue) according to the behavioral indicators during collaborative learning task. A hypothesis test was performed to explore the relationship between collaboration behavior and physiological synchrony during collaborative learning. At last, the classification analysis was applied to test the effectiveness of physiological synchrony as a predictor of collaborative learning quality.

### Define Collaborative and Independent Dyads

In the current study, two raters watched the videos of the group discussion and categorized the dyads into high or low collaboration style without being aware of the physiological data analysis. Effective interaction includes verbal and non-verbal interactions, such as tight conversation, eye contact, and joint attention on course material. The two raters recorded the time range of key interaction events in the video for each pair of students. For groups with 3 people, when the total time of interaction events exceeds 1/3 of the total time of PT, this pair is then categorized to collaborative dyads (CD), otherwise to independent dyads (ID). For the one group with 4 people, the threshold is set to 1/6 since there are six different pairs of students sharing the total interaction time. Fifteen dyads were sorted to CD and fourteen dyads to ID.

**Table 1.** Collaboration events and descriptions.

| Collaboration events | Description |
| --- | --- |
| Tight conversation | Discussion on one focused point between two people for several rounds |
| Eye contact | Though not talking, the person communicates with their teammates by eye contact, expressing agreement and disagreement on the topic |
| Joint attention | Two people show joint attention on the third object, such as course material, laptop screen, or the blackboard in the front of the classroom. |

### Engagement and Emotional Statement

It is also important to check the validity of experimental settings in naturalistic classroom. First, students’ engagement levels should be the same across the two sessions to ensure that any identified differences are not due to engagement differences. Second, students’ self-report on their subjective experience during IT and PT should be compared to check if the IT and PT did mean different learning strategies to them.

According to Figure 3, Engagement difference was not found between IT and PT. The participants were equally and highly engaged in both the independent task (*M* = 2.76, *SD* = 0.951) and group discussion task (*M* = 3.00, *SD* = 1.035), *t*(28) = 1.565, *p* = 0.258. This result suggests that participants were equally and highly engaged in both IT and PT, making engagement less likely to be the possible confounding factor for the additional synchrony during collaboration.

Same analysis was also conducted on the three affective dimensions. During group discussions, participants were more aroused (PT: *M* = 2.03, *SD* = 0.944; IT: *M* = 1.14, *SD* = 0.833, *t*(28) = 3.455, *p* = 0.002) and experienced higher positive emotion (PT: *M* = 2.93, *SD* = 0.813; IT: *M* = 2.11, *SD* = 0.875, *t*(27) = 5.037, *p* < 0.001). In the independent sessions, the participants reported to be more in control to their situation (PT: *M* = 1.90, *SD* = 0.673; IT: *M* = 2.79, *SD* = 1.013, *t*(28) = -5.363, *p* < 0.001). Higher score on arousal and valence indicated pleasant and excited discussion atmosphere during the collaborative learning task. Lower score in dominance is reasonable during PT since the process of multi-personal discussion came with negotiation and compromise (Figure 4).

**Figure 4.**
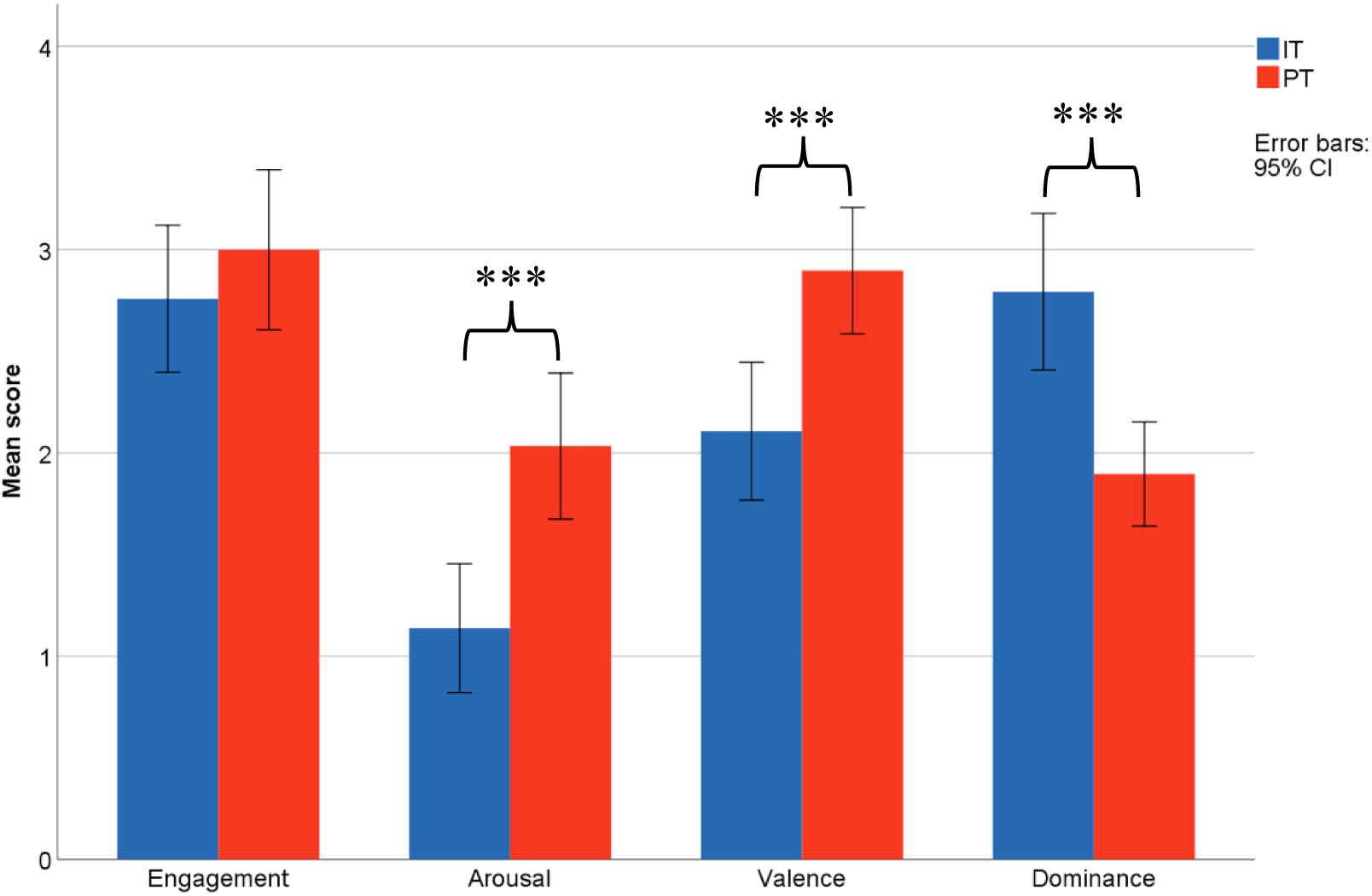
Engagement and emotional state between IT and PT \*\*\**p* < 0.001.

### Physiological Synchrony

The algorithm for the computation of GSR synchrony was adopted from Marci and Orr (2006), and calculated the moment-by-moment physiological concordance, named single session index (SSI). Same algorithm was also implemented on heart rate.

First, the 10Hz signal was further down sampled by averaging the ten numbers in each second. The moment-by-moment slope of the 1Hz data for each signal were then calculated using a 5-second window with a regression model at a 1-second roll-rate. Next, Pearson correlations were conducted on the slope for each pair of data with a 15 second window rolling at the rate of 1 second, reflecting a moment-by-moment synchrony in the last 15 seconds. The SSI is an index that shows the synchrony over a longer time period. It is the natural logarithm of the ratio of the sum of positive correlation coefficients divided by the absolute value of the sum of negative correlation coefficients over a given period of time. A sample of each step of the signal processing and synchronization calculation for IT and PT is shown in Figure 5.

**Figure 5.**
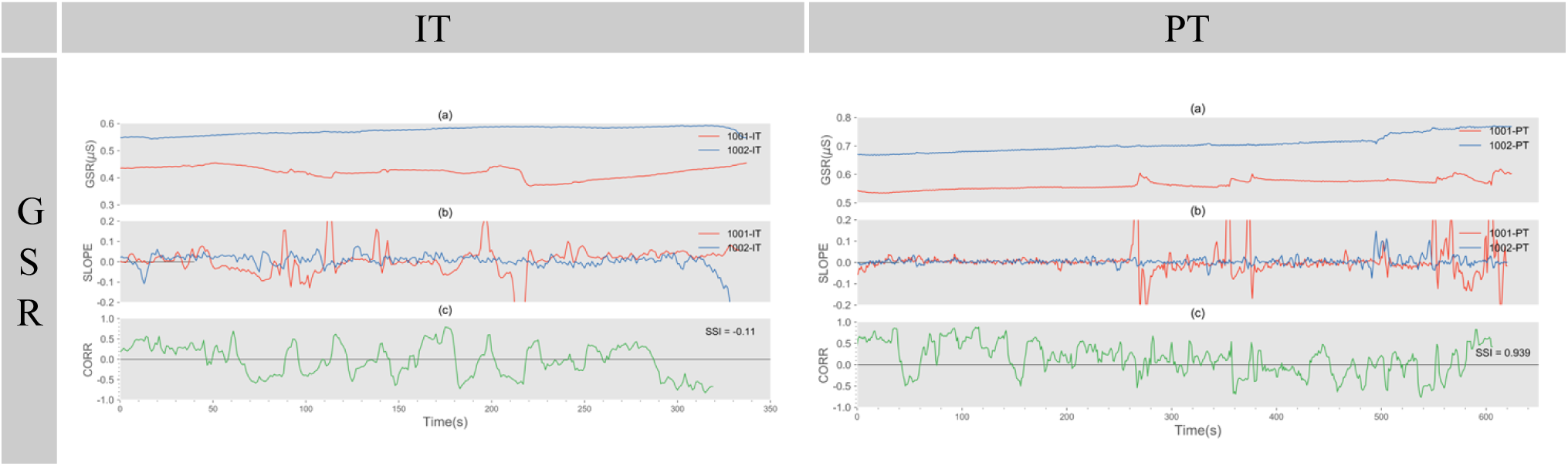
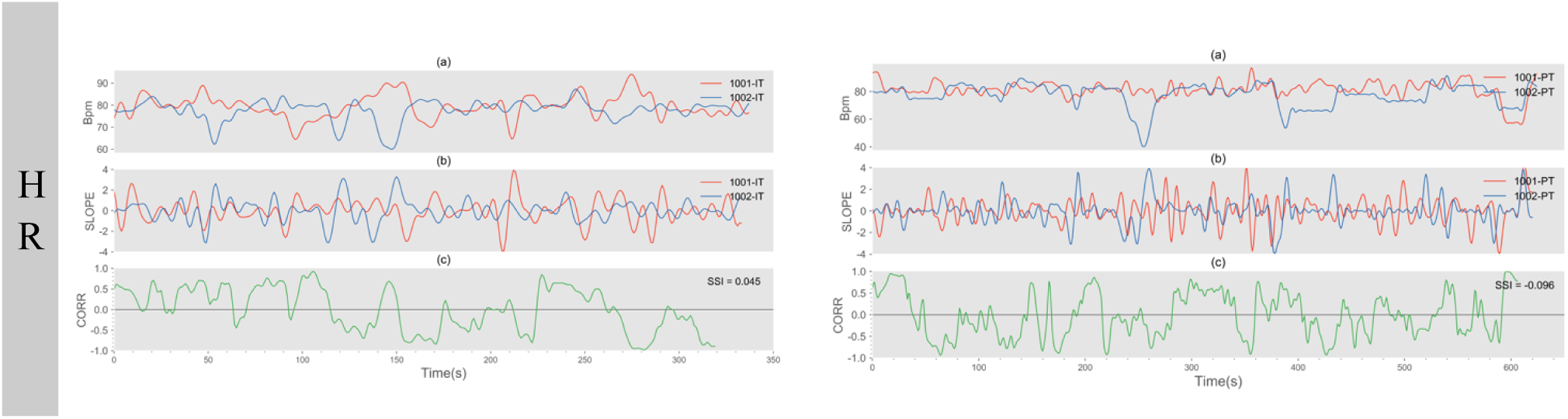
A sample of GSR and HR data processing.

In the upper left figure, (a) is the raw GSR signals of pair No. 10 in IT; (b) is the trajectory of slope for a 5-second window, rolling on the rate of 1 second; (c) shows the moment-by-moment correlation coefficients on a 15-second window. The three panels are the same as in the upper right figure. This is an example of the same pair in the collaborative task. The synchrony (SSI) was higher than that of in IT. The figures in the second row are examples of HR data.

It is interesting to find that synchrony on skin conductance during IT and PT reflects different styles of learners. When doing the independent task, ID (*M* = 0.322, *SD* = 0.449) showed similar synchrony level with their CD peers (*M* = 0.009, *SD* = 0.485), *t*(27) = 1.800, *p* = 0.084; while during the group discussion, the synchrony among ID (*M* = -0.170, *SD* = 0.396) was significantly lower than the CD (*M* = 0.231, *SD* = 0.380), *t*(27) = 2.781, *p* = 0.010 (Figure 6).

**Figure 6.**
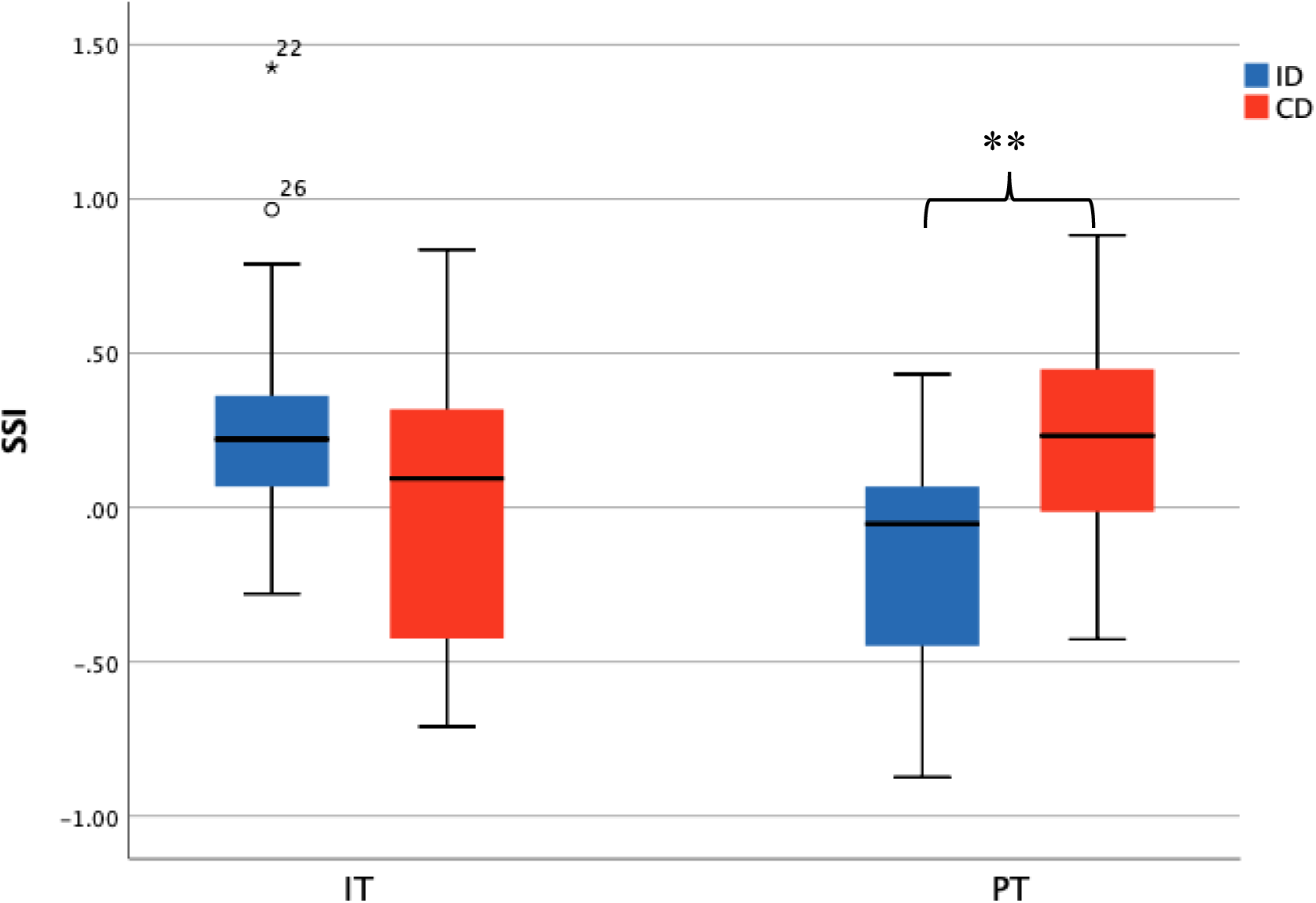
SSI among CD is significantly higher than that in ID during collaborative learning tasks. \*\**p* < .01.

Same analysis was conducted to explore the difference of synchrony of heart rate. When doing IT, ID (*M* = 0.171, *SD* = 0.462) and CD (*M* = -0.158, *SD* = 0.637) showed insignificant difference, *t*(26) = 1.157, *p* = 0.129; and during PT, there was also no significant difference between the synchrony among ID (*M* = -0.076, *SD* = 0.446) and the CD (*M* = -0.106, *SD* = 0.327), *t*(26) = 0.203, *p* = 0.841 (Figure 7).

**Figure 7.**
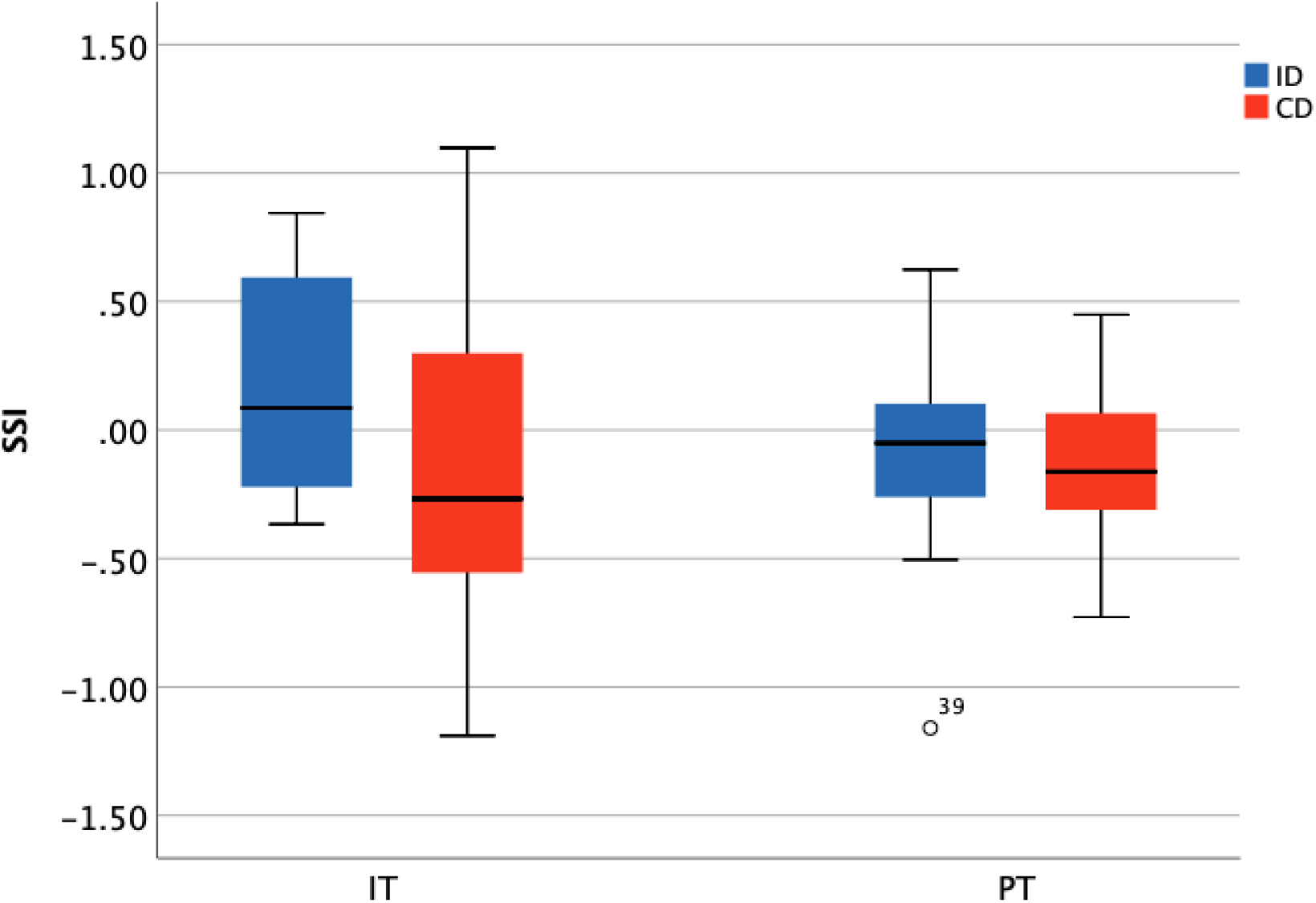
HR synchronization did not show significant difference between ID and CD during IT and PT.

### Physiological Synchrony as a Predictor of Collaborative Learning Quality

Since there is strong correlation between the interaction level and GSR synchrony, a ROC analysis was performed to test the accuracy of synchrony as a predictor of the collaborative learning quality in both IT and PT. Results showed that SSI is an acceptable predictor of collaboration quality for collaborative task (AUC = 0.767, p = 0.015). Synchrony did not discriminate different collaboration styles during IT (AUC = 0.343, p = 0.15) which is good since there was no collaborative behaviors and no significant difference in synchrony between CD and ID dyads during IT. The results of IT and PT together verified the robustness of synchrony as the predictor for collaborative learning quality. (See Figure 8.)

**Figure 8.**
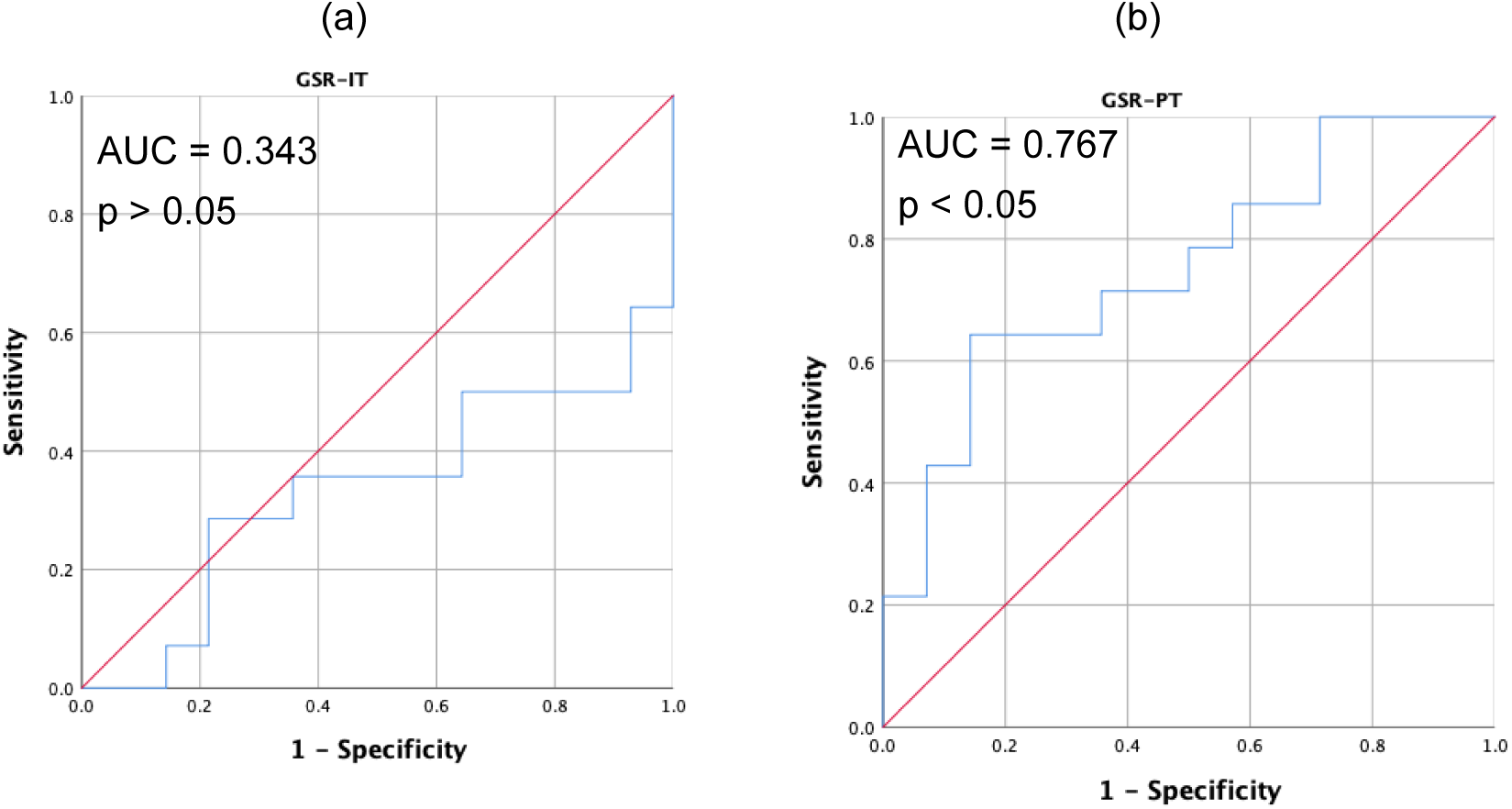
The ROC curve of GSR for IT (panel a) and PT (panel b) using synchrony as predictor for collaborative learning quality.

Same analysis was also applied on HR data. As shown in figure 9, the synchrony of HR exhibited low accuracy in predicting collaboration style (AUC = 0.454, p = 0.679) during PT. This is consistent with the low HR synchrony and undifferentiated HR synchronization level across collaborative learning quality.

**Figure 9.**
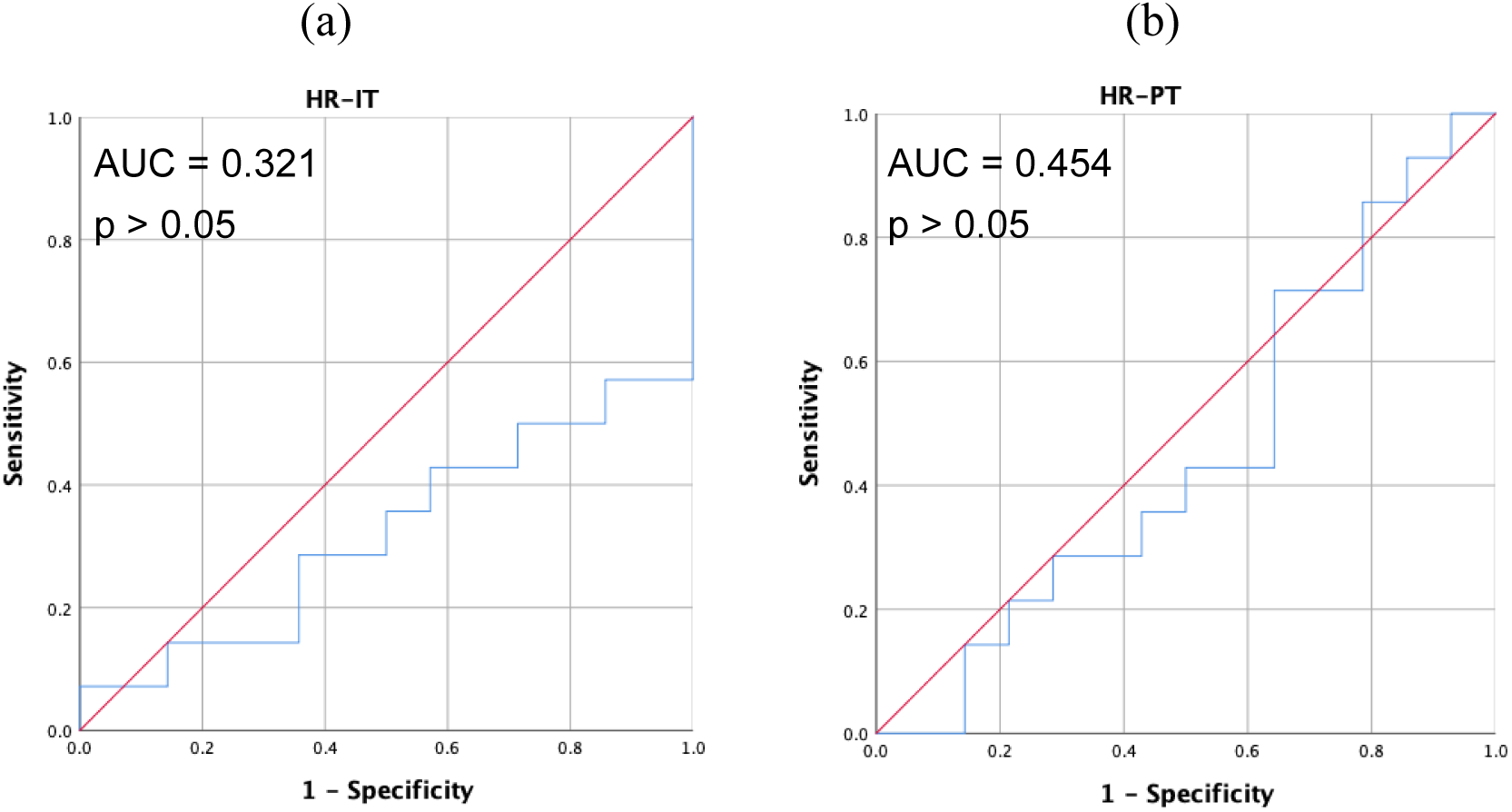
The ROC curve of HR for IT (panel a) and PT (panel b) using synchrony as predictor for collaborative learning quality.

## Conclusion and Discussion

The aim of the present study is to explore potentials of using physiological synchrony to measure collaboration quality in real educational settings, based on consistently identified synchrony during interpersonal interaction by previous studies. Existing studies show that learners are diverse in learning style and collaborative learning can manifest this diversity while students take different roles in the learning process (Pashler, McDaniel, Rohrer, & Bjork, 2008; Smith & Macgregor, 1992).

In the current study, the participants were categorized into collaborative dyads (CD) and independent dyads (ID) according to their natural behaviors in the collaborative learning task, and this behavioral difference significantly correlated with skin conductance synchrony. The results showed that participants who were categorized as CD during group discussions were associated with higher skin conductance synchrony. However, there was no significant difference in HR synchrony.

One possible explanation for this inconsistent results in skin conductance and heart rate may have to do with the fact that talking affects one’s cardiovascular system but not skin conductance. Talking, even without emotional expression, can increase the blood pressure of hypertension patients (Le Pailleur, Montgermont, Feder, Metzger, & Vacheron, 2001). Simple mental and verbal activities also affect heart rate variation through changes in respiratory frequency (Bernadi et al., 2000). On the other hand, no evidence was found for the correlation between EDA and free talking (Fowles, Kochanska, & Murray, 2000). In the natural group discussion context, students focused on the same task, trying to forge an integrated answer.

Among CD, two students were tuning their emotional state during the discussion, resulted in higher synchrony in EDA. But when two people talk, they talk in turns, not simultaneously. Actually, when participants doing verbal and motor activities in unison, HR synchrony was significantly higher than during unsynchronized moments (Müller & Lindenberger, 2011; Noy, Levit-Binun & Golland, 2015). Therefore, even the cognitive and emotional elements that generated synchrony in EDA may also synchronize HR in cognitive tasks as reported in the laboratory based studies (Henning & Korbelak, 2005; Mitkidis, McGraw, Roepstorff, & Wallot, 2015; Montague, Xu & Chiou, 2014), the effect could be mixed with that of talking on one’s HR. While on the other hand, EDA synchrony was identified during unstructured conversation (Silver & Parente, 2004).

Classification analysis proved that physiological synchrony may serve as a good indicator for interpersonal interaction quality. Higher physiological synchrony is positively correlated with higher interaction level. That is, higher frequency and longer time of interaction behaviors. Similar approach can be found in the research of predicting communication behavior using neural or physiological synchronization (Henning, Armstead, & Ferris, 2009; Jiang et al., 2012). This application can help identify different collaborative learning quality of the learners. It can also give instructors feedback on course content. One student may be attracted to one topic or interaction scheme but disinclined to another. In such case, physiological synchrony can provide clues in teaching adjustment.

Our findings provide evidence for the potential application of biosensors in the real-world classroom. We focus on the connection between the bio-signals and human behaviors on which we believe is the advantage of this interdisciplinary research area. This project also suggests that future researches in the same realm place attention to the scope of appropriate assumptions and research questions so that the laboratory-based experiments and naturalistic setting studies can be good complement for each other.

Students’ immediate learning outcome was not evaluated as the tasks were open-ended class discussions. Next, we will choose class sessions that has planned quizzes as a measure of learning performance.

## Acknowledgement

The authors appreciate Dr. Dan Zhang’s insightful comments on this study. This study is funded by the research grant of IFEE, Tsinghua University (2019IFEE003).

